# A multimodal precision oncology platform integrating genomics, single-cell transcriptomics, and functional drug screening expands therapeutic options in pediatric brain tumors

**DOI:** 10.64898/2026.10.12.758619

**Authors:** Xiaomeng Huang, Yi Qiao, Gage S. Black, Casey R. Sederman, Huiyi Yang, Teresa Musci, Alli Ludwick, Gongping He, Osama Youssef, Moises Gutierrez, Jingye Yang, Priya P. Chan, Trevor H. Memmott, Nicholas S. Whipple, Carol S. Bruggers, Aaron Quinlan, Philip J. Moos, Samuel H. Cheshier, Gabor T. Marth

## Abstract

Precision oncology for pediatric brain tumors mainly relies on clinically validated, targetable genomic alterations, which are often sparse, yield few matched therapies, and do not guide chemotherapy selection. We developed a tiered framework in which deep whole-genome sequencing and single-cell RNA sequencing extend targeted-therapy selection, with ex vivo functional drug screening supporting selection of both targeted therapies and chemotherapies. We applied it to 17 children with malignant CNS tumors. The approach expanded recommendations beyond genomics: functional screening provided every patient with at least one high-confidence targeted-therapy or chemotherapy nomination, while single-cell RNA sequencing identified an average of 37 druggable gene targets per patient, producing a fourfold increase in high-confidence targeted-therapy recommendations. In an exploratory analysis, functional classifications were directionally consistent with outcomes in 8 of 10 evaluable cases, although combination treatments and radiation precluded causal attribution. Single-cell profiling also identified subclone-specific vulnerabilities, supporting prospective evaluation of the platform’s feasibility and predictive utility.

## INTRODUCTION

Pediatric central nervous system (CNS) tumors are the most common solid tumors in children and the leading cause of cancer-related mortality in this population^1^. Survival varies markedly by tumor type and grade. For example, five-year survival is approximately 50% or lower for Group 3 medulloblastoma^2^ and below 20% for pediatric high-grade glioma^3^. Patients with aggressive tumors may be refractory to standard therapy or experience relapse, after which effective treatment options and evidence-based guidance are limited. Personalized approaches that use molecular tumor features to inform therapy are therefore needed^4^.

Over the past decade, several large studies have demonstrated the feasibility and potential clinical value of genomic profiling in children with high-risk or relapsed cancers (Table 1). TRICEPS identified potentially actionable alterations in 87% of patients using whole-exome and RNA sequencing^5^; the INFORM registry associated matched therapy for very-high-priority targets with longer progression-free survival^6^; and MAPPYACTS and iTHER reported matched therapy in 14-30% of patients, with responses in subsets of treated patients^7,8^. ZERO/PRISM reported two-year progression-free survival of 26% among patients receiving molecularly guided treatment versus 12% among those receiving standard care^9^; because treatment assignment was not randomized, this comparison should not be interpreted as proof of causal benefit. NCI-COG Pediatric MATCH identified predefined actionable alterations in 31% of 1,000 screened tumors^10^, whereas the SickKids KiCS program reported clinically actionable variants in 56% of 300 patients using integrated somatic-germline sequencing^11^. Collectively, these programs^12^ show that broader sequencing detects oncogenic events missed by targeted panels. Nevertheless, many alterations remain undruggable, and only a subset of patients receive matched therapy, underscoring the limitations of mutation-centric precision oncology.

**Table 1.** Comparing to existing pediatric precision oncology platforms.

| Platform | Genomics | RNA<br>- fusion | RNA<br>- expression | Functional<br>screen | Subclonal<br>resolution | Prioritization<br>framework<br>beside MTB |
| --- | --- | --- | --- | --- | --- | --- |
| Pediatric<br>MATCH | Targeted<br>DNA panel | Panel-based<br>RNA fusion | - | - | - | Predefined<br>actionable<br>alteration list |
| KiCS<br>(SickKids) | WES/WGS<br>+ germline | Bulk RNA-seq | - | - | - | Actionable<br>alteration list |
| ZERO /<br>PRISM | WGS | Bulk RNA-seq | Bulk outlier | - | - | Tiered priority<br>levels |
| MAPPYACTS | WES +<br>cfDNA WES | Bulk RNA-seq | - | - | - | “Ready for<br>routine use”<br>alteration list -><br>alteration-drug<br>match |
| iTHER | WES + low-<br>coverage<br>WGS | Bulk RNA-seq | Bulk outlier | - | - | INFORM<br>prioritization<br>method |
| INFORM | WES + low-<br>coverage<br>WGS | Bulk RNA-seq | Bulk outlier | - | - | Priority levels 1-7 |
| INFORM<br>DSP pilot | (inherits<br>registry<br>omics) | - | - | Yes (~70%<br>success) | - | Functional layer,<br>reported<br>separately |
| Treehouse<br>CARE | - | - | Bulk outlier<br>(sole<br>modality) | - | - | Outlier ranking |
| FIU FPM | UCSF500 | - | - | Yes (~84%;<br>expanded<br>cultures) | - | Functional-first |
| PNOC008 | UCSF500 +<br>WES +<br>WGS | Bulk RNA-seq | Not specified | - | - | Actionable<br>alteration list;<br>CNS-TAP (drug<br>recommendation<br>tool from<br>mutations) |
| **This platform** | **Deep WGS**<br>**120×** | **Bulk RNA-seq** | **scRNA-seq,**<br>**tumor-cell-**<br>**resolved** | **Yes (94%;**<br>**unexpande**<br>**d primary**<br>**cells)** | **Yes**<br>**(scBayes)** | **Two-track T/C**<br>**tiering** |

Complementary approaches have been proposed. RNA expression frameworks, such as the Comparative Analysis of RNA Expression (CARE) from the UC Santa Cruz Treehouse Initiative, use bulk RNA-seq outlier analysis to nominate overexpressed targets and have reported findings of potential clinical significance in 94% of pediatric patients^13,14^. A large clinical RNA-sequencing study from SickKids reported actionable findings in 87% of more than 2,300 tumors^15^. Ex vivo drug testing may further expand candidate options: a Florida International University functional precision medicine study reported improved progression-free survival in 83% of pediatric patients who received functionally guided treatment^16^, and the INFORM Drug Sensitivity Profiling pilot^17^ and other studies^18,19^ demonstrated operational feasibility. Most reported transcriptomic nomination frameworks use bulk RNA sequencing, which cannot directly separate tumor-cell expression from stromal and immune signals - a relevant limitation when tumor-cell content varies widely. To our knowledge, systematic integration of deep genomic sequencing, cell-resolved transcriptomics, and functional screening within one pediatric CNS tumor therapy prioritization framework has not been extensively evaluated.

Here we report a precision-oncology platform for pediatric brain tumors that integrates deep whole-genome sequencing (WGS), single-cell RNA sequencing (scRNA-seq), and ex vivo functional drug screening. Tumor WGS at an aggregate coverage of 120x was intended to improve detection of lower-frequency alterations relative to shallower sequencing, although sensitivity remains dependent on tumor purity, copy number, and the variant-calling threshold. scRNA-seq provided cell-resolved expression profiles that could distinguish malignant-cell signals from surrounding intrinsic stromal (e.g. astrocytes, oligodendrocyte), extrinsic stromal (e.g. fibroblasts, endothelial cells), and immune-cell populations (e.g. T cells, macrophages etc.) and examine intratumoral heterogeneity^20–22^. Ex vivo screening supplied an independent, cohort-relative measure of drug response that cannot be inferred from molecular data alone. We integrated these layers in a transparent framework for prioritizing candidate therapies according to the available evidence. We applied the strategy to 17 children with CNS tumors to evaluate feasibility, therapy nomination yield, and the resolution added by each modality.

## RESULTS

### Study Design and Patient Enrollment

This feasibility study was conducted under protocols approved by the University of Utah Institutional Review Board (IRB #00121939 and IRB #00112349). We obtained tissue (tumor and peripheral blood) from seventeen patients with diagnosed malignant central nervous system (CNS) tumors after obtaining written informed consent. All patients underwent surgical resection of their primary tumors at Intermountain Health Primary Children’s Hospital. Although precision oncology has conventionally been reserved for relapsed or recurrent disease, our aim is to apply it as early as clinically actionable - realistically at post-induction maintenance - to forestall recurrence in patients at high risk, rather than only to treat it once established. Tissue availability reinforces this timing: comprehensive profiling requires viable surgical tissue, which is reliably obtained at primary resection, whereas recurrent tumors are often unresectable. When a relapse cannot itself be profiled, the primary specimen provides the molecular basis for second-line decisions. Profiling at diagnosis therefore positions the platform to guide both maintenance and, where feasible, therapy selection at the time of relapse.

The cohort was diagnostically diverse and included atypical teratoid/rhabdoid tumor (ATRT; n=2), embryonal tumor with multilayered rosettes (ETMR; n=1), pediatric-type diffuse high-grade glioma [PDHGG; H3 wild type and IDH wild type (n=3), infant-type hemispheric glioma (n=1)], medulloblastoma [MB; SHH-driven (n=1), Group 3 (n=4), and Group 4 (n=4)], and posterior fossa ependymoma group PFA (EPEN; n=1) (Figure 1C). All tumor diagnoses were verified by a board-certified neuropathologist. Molecular profiling—incorporating the UCSF 500 targeted gene sequencing panel^23^ and genome-wide DNA methylation array classification^24^ -was conducted in a CLIA-certified, CAP-accredited clinical laboratory in accordance with institutional diagnostic standards. Clinical and demographic characteristics are provided in Supplemental Table 1.

**Figure 1.**
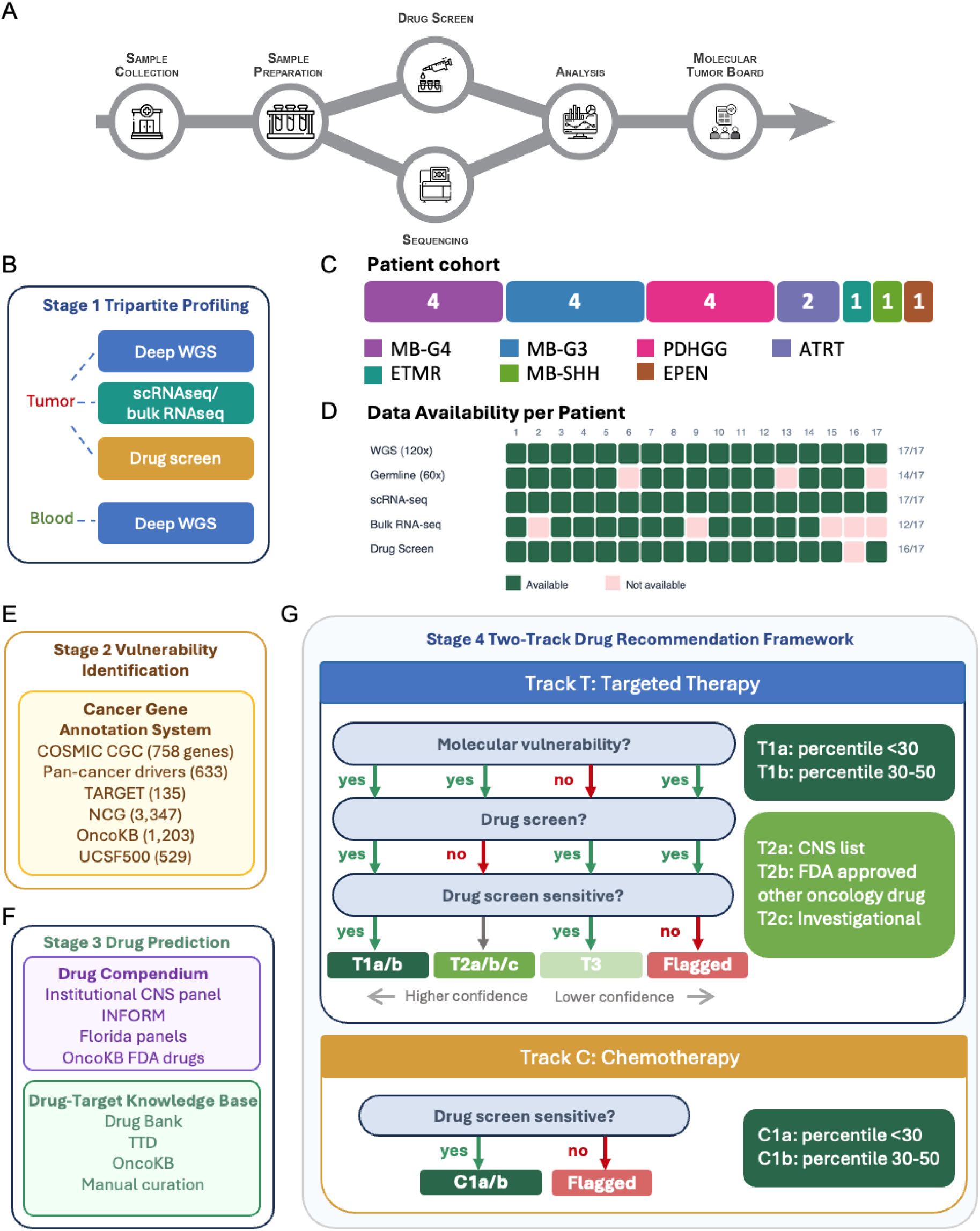
Overview of the multimodal precision oncology platform and integrated drug-recommendation framework. **(A)** End-to-end clinical workflow, proceeding from surgical sample collection and sample preparation through parallel ex vivo drug screening and next-generation sequencing, followed by integrated analysis and review at a molecular tumor board. **(B)** Stage 1, tripartite molecular and functional profiling. Fresh tumor tissue is divided into three parallel workflows: deep WGS, single-cell and bulk RNA sequencing, and ex vivo drug screening; with matched peripheral blood used for germline WGS. **(C)** Composition of the 17-patient cohort by histologic diagnosis: medulloblastoma Group 3 (MB-G3, n=4), medulloblastoma Group 4 (MB-G4, n=4), pediatric-type diffuse high-grade gliomas (PDHGG, n=4), atypical teratoid/rhabdoid tumor (ATRT, n=2), embryonal tumor with multilayered rosettes (ETMR, n=1), medulloblastoma SHH (MB-SHH, n=1), and ependymoma (EPEN, n=1). **(D)** Data availability across the five assays for each of the 17 patients. Filled cells indicate available data. **(E)** Stage 2, vulnerability identification. Somatic mutations and differentially expressed genes are annotated against a cancer gene annotation system integrating six curated resources. **(F)** Stage 3, drug prediction. Candidate therapies are drawn from a multi-source drug compendium (institutional pediatric CNS panel, INFORM and Florida functional-medicine panels, and OncoKB FDA-approved drugs) and mapped to vulnerabilities through a drug-target knowledge base (DrugBank, TTD, OncoKB, and manual curation). **(G)** Stage 4, two-track drug-recommendation framework. Track T (targeted therapy) assigns drugs to tiers by integrating molecular vulnerability, drug-screen availability, and ex vivo sensitivity: T1a/b (molecular-functional concordance; percentile <30 or 30-50), T2a/b/c (molecular only, not screened; CNS-relevant, FDA-approved, or investigational), T3 (functional only, percentile <50), and Flagged (molecular only, percentile ≥50). Track C (chemotherapy) relies solely on screen, assigning C1a/b (percentile <30 or 30-50) or Flagged (percentile ≥50).

### Precision oncology workflow including comprehensive omic-profiling, ex-vivo drug screening and integrated drug prioritization

Our platform proceeds through four sequential stages from tumor tissue to therapeutic recommendations (Figure 1A): (1) tripartite molecular and functional profiling of fresh surgical specimens; (2) identification of cancer-relevant vulnerabilities through a cancer gene annotation system; (3) translation of vulnerabilities into candidate therapeutics via a drug-target knowledge base and curated drug compendium; and (4) integrated prioritization through a two-track drug recommendation framework.

### Unique multimodal molecular and functional profiling

Fresh tumor tissue was processed through three parallel workflows (Figure 1B): (1) tumor WGS at a target coverage of 120x (17/17 patients), a higher-than-typical sequencing depth to help with the identification of subclonal tumor mutations, with matched germline sequencing at 60x for 14 patients and tumor-only analysis using a cohort-derived panel of normals for three patients; (2) scRNA-seq for cell-resolved expression profiling, supplemented by bulk RNA-seq in 12 patients for fusion detection; and (3) ex vivo screening with a curated 25-drug panel (16/17 patients) to measure short-term drug response in dissociated cultures. Assay availability by patient is shown in Figure 1D. To our knowledge, this is an early evaluation of deep WGS, single-cell transcriptomics, and functional screening within a unified pediatric CNS tumor nomination framework.

### Comprehensive cancer gene annotation system and drug-target knowledge base

We constructed a cancer-gene annotation set to enable the interpretation of cancer-relevant genomic and transcriptomic vulnerabilities, utilizing six data resources: the COSMIC Cancer Gene Census (758 genes)^25^, a pan-cancer driver-gene set (633 genes)^26^, TARGET^27^, the Network of Cancer Genes (3,347 genes)^28^, OncoKB (1,203 genes)^29,30^, and the UCSF500 panel (529 genes) (Figure 1E; Supplemental Table 2). Somatic alterations from WGS and overexpressed genes from scRNA-seq were annotated against this set to identify cancer-associated candidates. A separate drug-target knowledge base integrated DrugBank^31^, the Therapeutic Target Database^32^, the Genomics of Drug Sensitivity in Cancer resource^33^, OncoKB^29,30^, and manual curation to map candidates to drugs with reported mechanisms of action (Figure 1F; Supplemental Table 4). These annotations nominate hypotheses; they do not by themselves establish functional dependence or clinical actionability.

### Systematic curation of a multi-source drug compendium

The relevant set of therapies for the treatment of the pediatric brain cancer patients was assembled from four sources: (1) a physician-curated, pediatric CNS tumor-relevant drug list (Supplemental Table 3); (2) the INFORM Drug Sensitivity Profiling panel^17^; (3) the Florida functional precision medicine panel^16^; and (4) all FDA-approved oncology drugs from OncoKB (Figure 1F) including on-label and off-label drugs for the treatment of CNS tumors. Twenty-five drugs were selected for ex vivo screening. This panel represents a tradeoff between feasibility of screening within a clinically relevant timeframe, clinical relevance to pediatric CNS tumors, and mechanistic coverage of key pathways including RTK-MAPK, PI3K/AKT/mTOR, cell-cycle regulation, epigenetic modifiers, and cytotoxic agents (Supplemental Table 3).

Drug response was quantified by area under the dose-response curve (AUC). For each drug, a patient’s AUC was ranked against the available AUC values for that drug across the cohort (Figure 2A; Supplemental Figure 5). Lower percentiles indicate lower viability relative to other tested tumors for the same compound. This within-drug ranking reduces the influence of between-drug potency differences but does not define an absolute sensitivity threshold or establish clinical efficacy.

**Figure 2.**
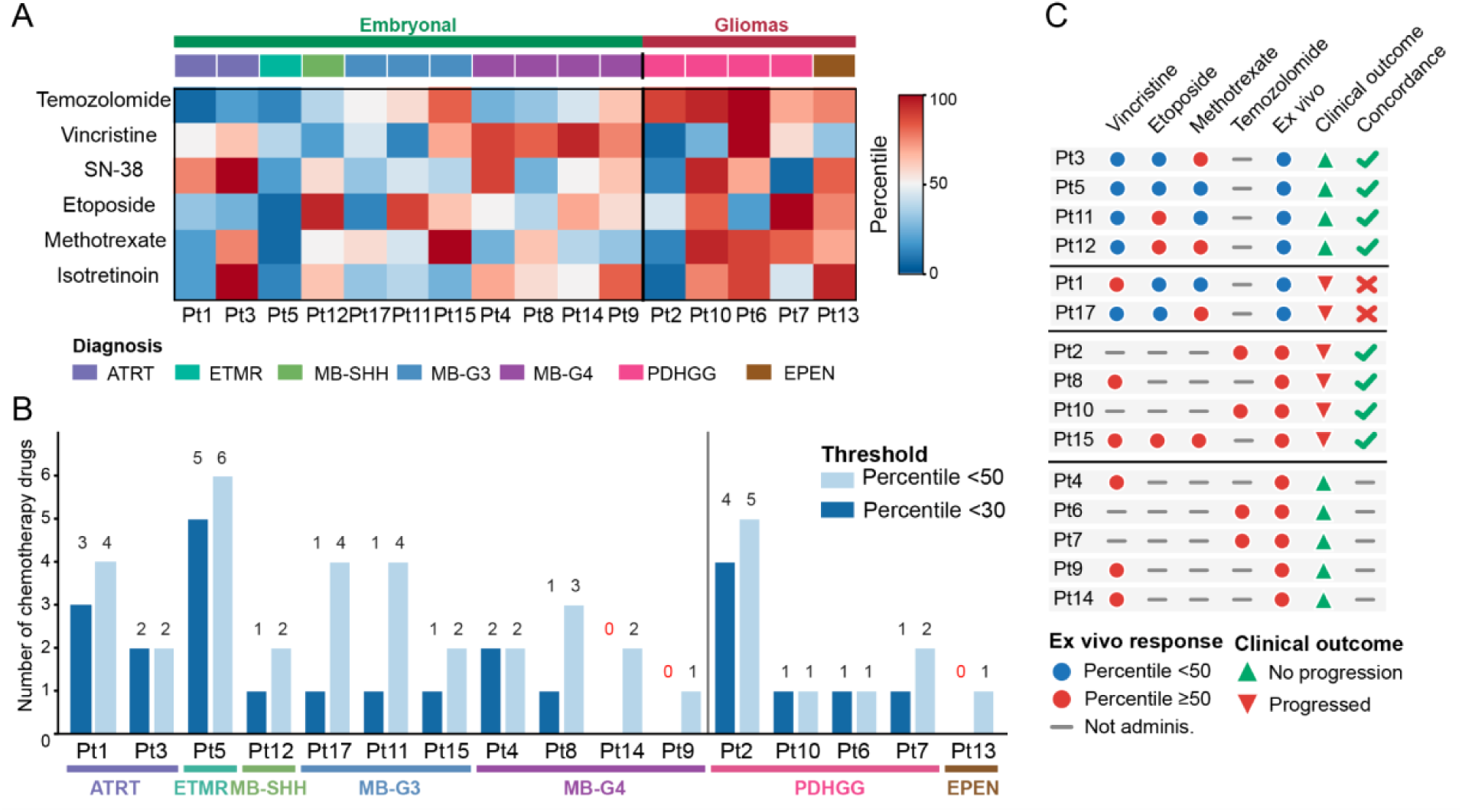
*Ex vivo* functional screening identifies cohort-relative response patterns and enables an exploratory comparison with clinical outcomes. (**A**) Heatmap of response percentiles across 16 screened patients for five cytotoxic agents (temozolomide, vincristine, SN-38, etoposide, and methotrexate) plus isotretinoin. Color indicates the within-drug cohort percentile, from lower AUC (blue) to higher AUC (red). Patients are grouped as embryonal tumors or gliomas; the color bar denotes histology. (**B**) Number of agents per patient below the 50th (light) and 30th (dark) percentiles. (**C**) Descriptive comparison of administered screened agents and clinical outcome for vincristine, etoposide, methotrexate, and temozolomide. Categories are an administered agent below the cohort median without relapse (n=4), an administered agent below the median with relapse (n=2), no administered agent below the median with relapse (n=4), and no administered agent below the median without relapse (n=5). Symbols indicate screen classification, administration, and outcome.

### Two-track therapy recommendation framework integrating molecular and functional evidence

We developed a two-track framework that separates the inference rules for targeted agents (Track T) and cytotoxic chemotherapy (Track C) (Figure 1G). In Track T, genomic alterations or transcriptomic outliers nominate drug-target hypotheses. When a nominated drug was included in the ex vivo panel, cohort-relative response supplied an orthogonal line of evidence. Drugs with a matched molecular candidate and an AUC percentile below 50 were assigned T1: T1a for percentile <30 and T1b for percentile 30-50. T1 therefore denotes molecular-functional concordance within this framework. Molecularly nominated drugs not screened were assigned T2 and stratified as T2a (physician-curated pediatric CNS list), T2b (other FDA-approved oncology agents), or T2c (other investigational pediatric oncology agents in clinical trials). Screened targeted agents below the cohort median without a matched molecular candidate were assigned T3. A molecularly nominated drug with an AUC percentile ≥50 was flagged as discordant rather than labeled clinically resistant.

In Track C, the screen was the sole evidence source because the cytotoxic agents were not mapped to patient-specific molecular markers. Track C assignments were C1a (percentile <30), C1b (percentile 30-50), or Flagged-C (percentile ≥50). All percentiles are relative to the tested cohort and require prospective validation.

### Functional drug screening identifies patient-specific chemotherapy options for all screened patients

We screened dissociated tumor specimens against a curated 25-drug panel selected for pediatric CNS relevance (Supplemental Table 3; Methods). Cells were assayed directly (P0) or after one passage (P1), without extended culture. Sixteen of 17 specimens (94%) produced screening data (Figure 1D). This completion rate is comparable with rates reported by the INFORM Drug Sensitivity Profiling pilot (∼70%) and De La Rocha et al. (∼84%), although differences in eligibility, culture, assay, and success definitions preclude a direct performance comparison.

Under Track C, each of the 16 screened tumors had at least one of six agents below the cohort median (C1; median, 2 agents per patient; range, 1-6), and 12 of 16 (75%) had at least two. Thirteen patients (81%) had at least one agent below the 30th percentile (C1a; median, 1; range, 0-5), and 12 (75%) had at least one agent between the 30th and 50th percentiles (C1b; median, 1; range, 0-3) (Figure 2A,B). Fifteen tumors (94%) had at least one agent at or above the cohort median (Flagged-C; median, 4; range, 0-5). These designations represent relative rankings within this small, heterogeneous cohort and should not be interpreted as absolute clinical sensitivity or resistance. Four patients (Pt2, Pt8, Pt10, and Pt15) received regimens containing a screened agent classified as Flagged-C and later relapsed (Figure 2C); because treatment was multimodal, this descriptive association cannot identify the contribution of the individual agent or of functional screen-nominated alternatives. In an exploratory analysis, we compared cohort-relative functional screen classifications with subsequent clinical outcomes (Figure 2C). Pt13 received radiation alone and was not evaluable. The other 15 screened patients received at least one agent represented in the panel and were grouped descriptively as follows: an administered agent below the cohort median without observed relapse (Pt3, Pt5, Pt11, and Pt12); an administered agent below the median with relapse (Pt1 and Pt17); no administered agent below the median with relapse (Pt2, Pt8, Pt10, and Pt15); or no administered agent below the median without observed relapse (Pt4, Pt6, Pt7, Pt9, and Pt14).

This comparison is strongly confounded: all patients received radiation, systemic regimens contained multiple agents, only some administered agents were screened, and follow-up duration may differ. The first and third groups (8 patients) were directionally consistent with the screen classification, whereas the second group (2 patients) was not. The fourth group (5 patients) was not interpretable because favorable outcomes could reflect radiation or unscreened therapies; the same uncertainty limits attribution in the first group. These data therefore do not estimate predictive accuracy. A prospective study with prespecified thresholds, standardized outcome windows, and treatment assignment informed by the assay is required to evaluate predictive utility.

### Integration of genomic and transcriptomic vulnerabilities with functional evidence expands targeted therapy nominations

In our framework, targeted agents were first nominated from genomic alterations or transcriptomic outliers linked to reported pharmacological targets. The drug screen then supplied concordant or discordant functional evidence when the agent was included in the panel. We therefore characterized the genomic and transcriptomic candidate landscape and integrated it with ex vivo response to generate tiered nominations.

### Genomic vulnerabilities are sparse

Deep WGS was performed at a mean tumor coverage of 120x in all 17 patients; matched germline WGS at 60x was available for 14. With one exception, no pathogenic or likely pathogenic germline variants were reported in established childhood cancer-predisposition genes^34^. Pt10, carried a germline deletion of *MSH2* exons 1-3, consistent with this patient’s clinical diagnosis of Lynch syndrome. This patient also had a somatic *MSH2* stop-gain variant (p.Q419*) likely explaining the exceptionally high tumor mutation burden. Somatic SNVs and small insertions/deletions were predominantly private and nonrecurrent. Pt10 was a hypermutated outlier with 83,742 variants. Across the other tumors, short variants affected 1-30 cancer-associated genes and yielded 0-2 candidate targetable events; Pt10 had 196 and 15, respectively (Supplemental Tables 5 and 6). Copy-number alterations (Supplemental Figure 2) affected a reported mean of 302 annotated cancer-associated genes per patient (range, 0-1,398), of which 6.5 (range, 0-34) met the study’s targetability rules (Supplemental Table 5 and 6). Overall, the framework identified a reported mean of approximately 7.5 candidate genomic vulnerabilities per patient, motivating evaluation of complementary modalities.

### Single-cell transcriptomics reveals additional candidate tumor vulnerabilities

We performed scRNA-seq on all 17 tumors and bulk RNA-seq on 12 tumors for fusion detection. Tumor-cell content varied from 4% to 97%, illustrating the potential for bulk profiles to mix malignant and nonmalignant signals (Discussion). After quality control, 962-16,028 cells per sample were retained (110,373 total), and malignant cells were distinguished from stromal and immune populations using scBayes^35^ and inferCNV^36^ (Methods; Supplemental Figure 3). Comparing each patient’s malignant cells with a pan-cohort malignant-cell reference identified a mean of 37 and median of 42 cancer-associated, druggable overexpressed genes per patient (range, 2-77) (Supplemental Table 5). Candidates involved growth-factor/MAPK signaling (*ERBB4*, *MAP2K2*), cell-cycle control (*CDK4*), epigenetic regulation (*HDAC3*, *IDH1*), invasion (*MMP2*, *VEGFB*), and proteostasis or nuclear transport (*PSMB1/5*, *XPO1*). These patterns nominate expression hypotheses but do not establish functional dependence. Bulk RNA-seq also identified an *ALK* fusion in Pt7, with concordant malignant-cell *ALK* overexpression by scRNA-seq.

### Integrated targeted therapy recommendations across tiers

Integration yielded 48 T1 nominations across 16 screened patients (median, 2 per patient; mean, 3.0; range, 0-9); 13 patients (81%) had at least one T1 nomination (Figure 3A; Supplemental Table 7). By definition, T1 required a matched molecular candidate and an ex vivo AUC percentile below 50. These nominations therefore represent molecular-functional concordance within the framework.

**Figure 3.**
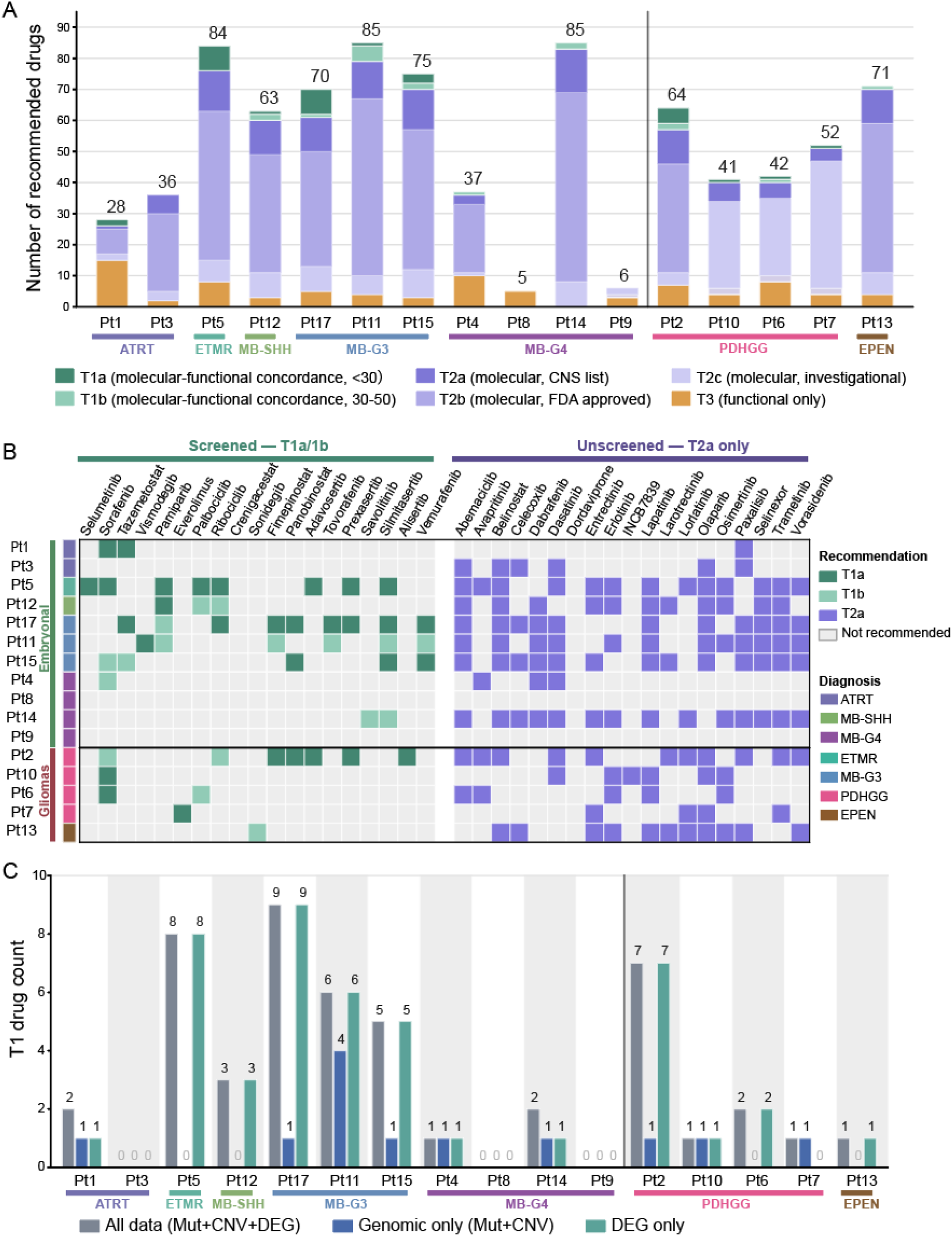
Multimodal integration increases the confidence of targeted-therapy recommendations and expands the therapeutic landscape beyond genomics alone. **(A)** Stacked bar chart of the total number of recommended drugs per patient, colored by tier (T1a, T1b, T2a, T2b, T2c, T3) and grouped by diagnosis. Total counts are shown above each bar. **(B)** Drug-by-patient matrix of tier assignments. Columns are drugs, split into agents included in the screening panel (T1) and agents not screened (T2a); rows are patients grouped into embryonal tumors and gliomas. Cell color indicates tier: T1a; T1b; T2a or not recommended. **(C)** Modality contribution analysis of high-confidence (T1) nominations. For each patient, T1 drug counts are recomputed using all data (mutations + CNV + DEG), genomic data only (mutations + CNV), or transcriptomic data only (DEG), quantifying the contribution of single-cell transcriptomics to high-confidence recommendations.

T2 captured molecularly nominated drugs that were not included in the 25-drug screen. Patients had a median of 8.5 T2a CNS-relevant candidates (Figure 3A, B), while T2b included a median of 36 FDA-approved oncology drugs and T2c included investigational agents. These large counts are nomination sets that require clinical filtering for evidence level, pediatric dosing, CNS penetration, toxicity, availability, and disease context. T3 included a median of 4 agents below the cohort median without a matched molecular candidate. A median of 2.5 Flagged-T agents per patient had a matched molecular candidate but an AUC percentile at or above 50, indicating molecular-functional discordance rather than proven clinical resistance.

Three patients had no T1 nomination. Pt3 (ATRT) had molecular candidates, but none of the corresponding screened agents had an AUC percentile below 50. Pt8 and Pt9 (Group 4 medulloblastoma) had agents below the cohort median but no matched molecular candidate under the study’s annotation rules. These patterns may reflect assay limitations, cohort-relative thresholds, incomplete annotation, or biology not captured by the framework; or possibly intrinsic resistance mechanisms.

We next quantified the respective contributions of genomic vs. transcriptomic vulnerability data to T1 assignments. With genomic candidates alone, the mean T1 count decreased from 3.0 to 0.75 per patient (75%); 9 of 16 patients lost at least half of their T1 nominations, and 4 had none (Figure 3C; Table 2). With transcriptomic candidates alone, the mean was 2.81 (6% lower than the all-data count); 3 patients lost at least half of their nominations, and 1 had none. Under this framework and annotation set, transcriptomic outliers generated most T1 nominations, while genomic alterations contributed a smaller number not recovered by transcriptomics alone.

**Table 2.**
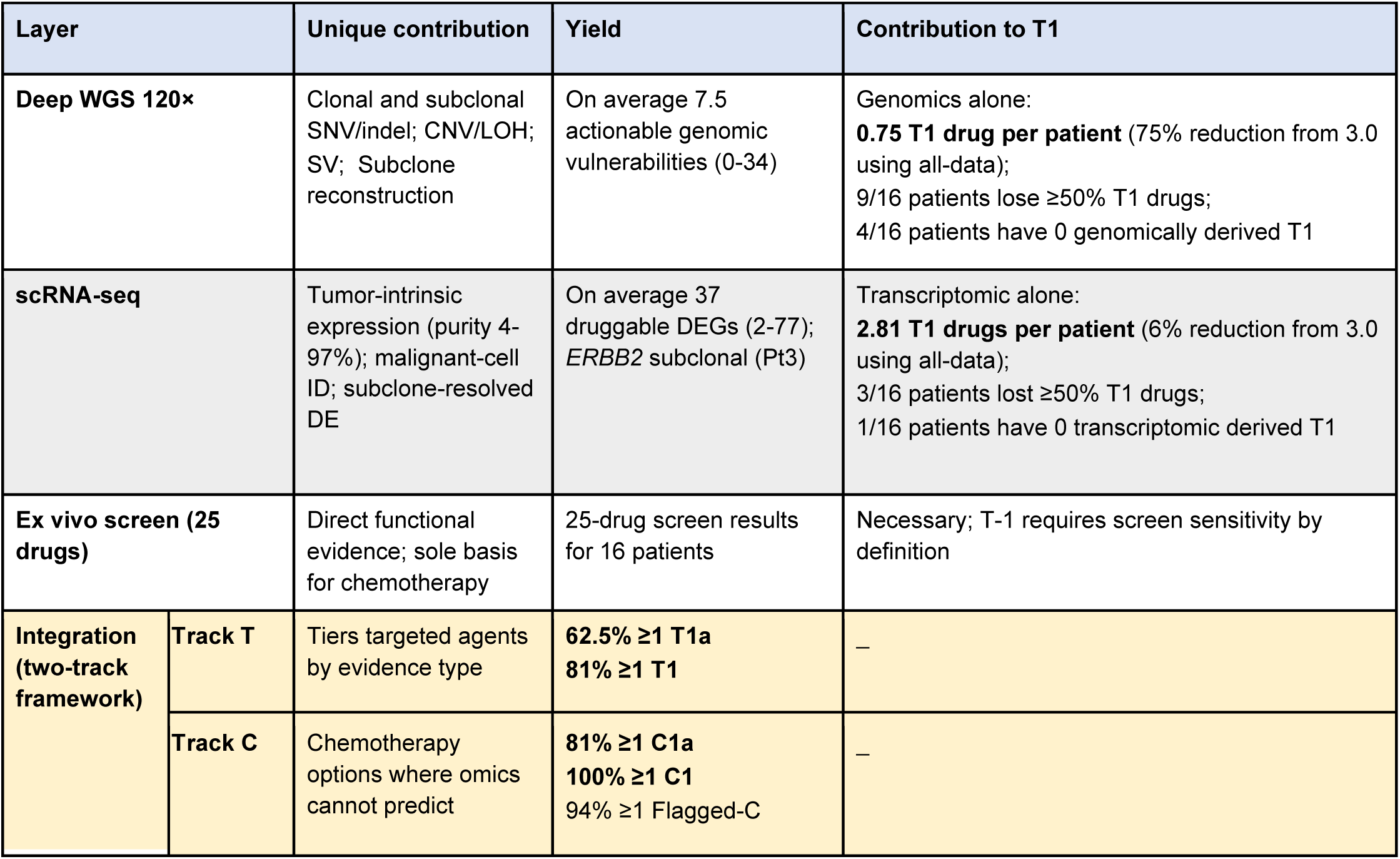
Contribution of each modality.

| Layer |  | Unique contribution | Yield | Contribution to T1 |
| --- | --- | --- | --- | --- |
| Deep WGS 120× |  | Clonal and subclonal SNV/indel; CNV/LOH; SV; Subclone reconstruction | On average 7.5 actionable genomic vulnerabilities (0-34) | Genomics alone:<br>**0.75 T1 drug per patient** (75% reduction from 3.0 using all-data);<br>9/16 patients lose ≥50% T1 drugs;<br>4/16 patients have 0 genomically derived T1 |
| scRNA-seq |  | Tumor-intrinsic expression (purity 4-97%); malignant-cell ID; subclone-resolved DE | On average 37 druggable DEGs (2-77); *ERBB2* subclonal (Pt3) | Transcriptomic alone:<br>**2.81 T1 drugs per patient** (6% reduction from 3.0 using all-data);<br>3/16 patients lost ≥50% T1 drugs;<br>1/16 patients have 0 transcriptomic derived T1 |
| Ex vivo screen (25 drugs) |  | Direct functional evidence; sole basis for chemotherapy | 25-drug screen results for 16 patients | Necessary; T-1 requires screen sensitivity by definition |
| Integration (two-track framework) | Track T | Tiers targeted agents by evidence type | **62.5% ≥1 T1a**<br>**81% ≥1 T1** | — |
|  | Track C | Chemotherapy options where omics cannot predict | **81% ≥1 C1a**<br>**100% ≥1 C1**<br>94% ≥1 Flagged-C | — |

### Illustrative cases demonstrate the clinical relevance of multimodal integration

We highlight two cases to illustrate how genomic, transcriptomic, and functional evidence contributed nonredundant information to patient-level nominations.

#### Pt2 (PDHGG)

This case illustrates that the three data modalities contribute non-redundantly, with concordant nominations, transcriptome-only nominations, and a functional screen-only negative finding all arising within a single tumor.

Two T1b nominations combined evidence from all three modalities. For sorafenib, a *KIT*/*PDGFRA* high-level copy gain (Figure 4A), malignant-cell overexpression of *KIT* and *RET* (Figure 4B), and a sorafenib AUC between the 30th and 50th cohort percentiles (Figure 4C) implicated receptor-tyrosine-kinase signaling. For ribociclib, homozygous *CDKN2A*/*CDKN2B* deletion, *CDK4* overexpression, and an AUC in the same percentile interval implicated cell-cycle signaling. The agreement represents molecular-functional concordance under the study’s tier rules; it does not validate target dependence or clinical efficacy.

**Figure 4.**
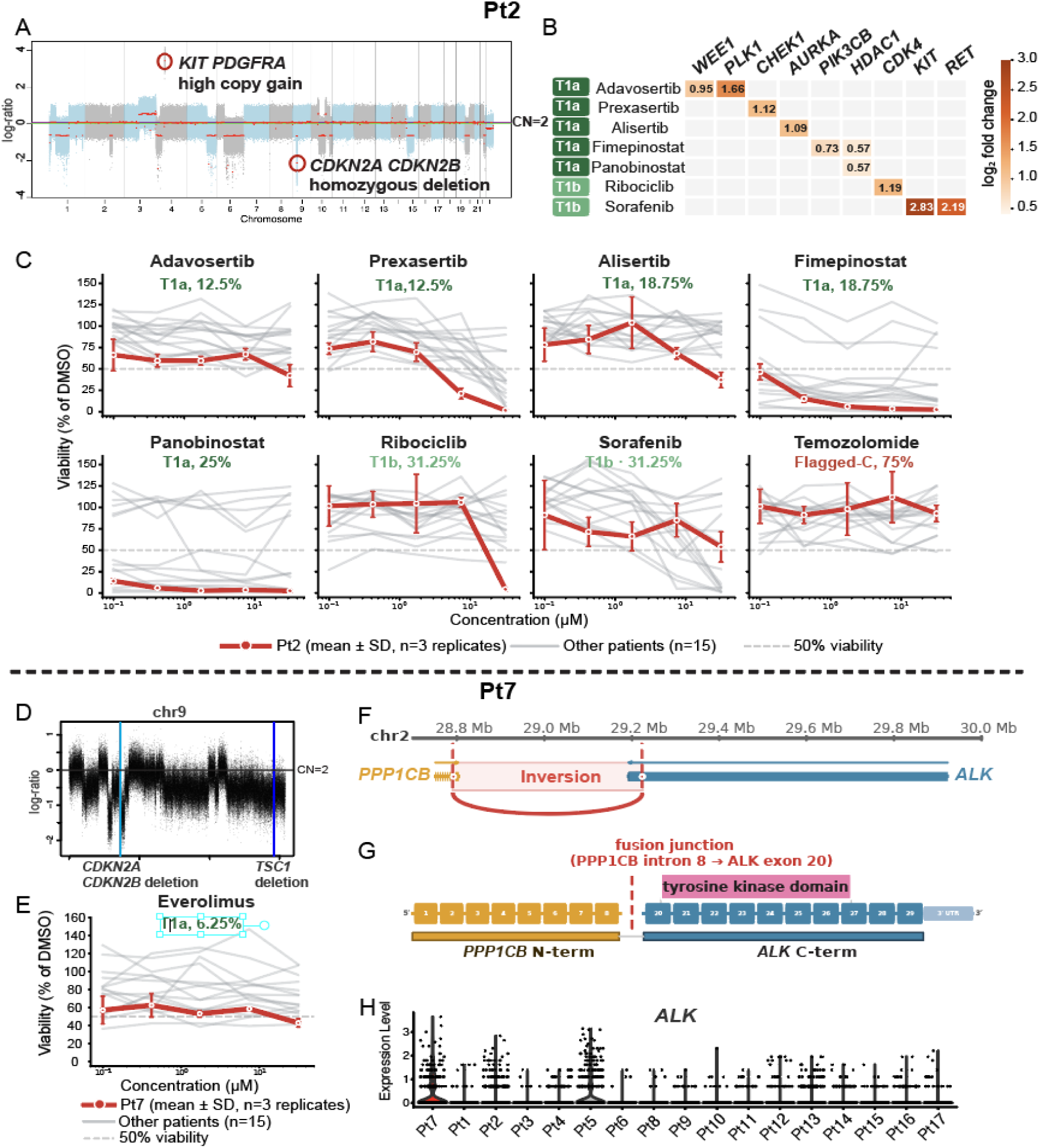
Illustrative cases demonstrating multimodal integration of genomic, transcriptomic, and functional evidence. (A-C) Patient 2. **(A)** Genome-wide copy-number profile (log ratio) showing a high-level *KIT*/*PDGFRA* copy gain and a homozygous *CDKN2A*/*CDKN2B* deletion. **(B)** Heatmap of targetable differentially expressed genes (tumor-cell log2 fold change) linking overexpressed targets - *WEE1*, *PLK1*, *CHEK1*, *AURKA*, *PIK3CB*, *HDAC1*, *CDK4*, *KIT*, *RET* - to recommended agents. **(C)** Ex vivo dose-response curves for eight drugs (Pt2 in red, mean ± SD of three replicates; other patients in gray), annotated by tier and percentile; temozolomide is flagged as screen-resistant (Flagged-C, percentile 75). (D-H) Patient 7. **(D)** Copy-number profile of chromosome 9 showing *CDKN2A*/*CDKN2B* and *TSC1* deletions. **(E)** Everolimus dose-response curve, indicating strong mTOR-inhibitor sensitivity (T1a). **(F)** WGS-derived structural variant: a chromosome 2 inversion juxtaposing *PPP1CB* and *ALK*. **(G)** Resulting in-frame *PPP1CB*-*ALK* fusion transcript detected by bulk RNA-seq (*PPP1CB* intron 8 to *ALK* exon 20), retaining the *ALK* tyrosine kinase domain. **(H)** scRNA-seq violin plots of *ALK* expression across all patients, showing *ALK* overexpression confined to Pt7 tumor cells.

Five additional T1a nominations involved DNA-damage or mitotic-checkpoint candidates (adavosertib-*WEE1*/*PLK1*, prexasertib-*CHEK1*, and alisertib-*AURKA*) and epigenetic-regulation candidates (fimepinostat-*HDAC1*/*PIK3CB* and panobinostat-*HDAC1*), each linked to an overexpressed gene and an AUC below the 30th cohort percentile (Figure 4B,C). Five of the seven T1 nominations in this tumor depended on transcriptomic candidates without a corresponding genomic lesion and therefore would not have been generated from WGS plus screening alone.

Conversely, temozolomide had an AUC at the 75th cohort percentile and was classified as Flagged-C (Figure 4C). The patient later progressed on a temozolomide-containing regimen. This temporal association is directionally consistent with the screen classification, but the combination regimen, radiation, cohort-relative threshold, and absence of a single-agent counterfactual preclude attributing progression to temozolomide or treating the result as evidence against standard of care.

#### Pt7 (PDHGG)

This case illustrates how genomic and transcriptomic evidence can nominate an agent that was not included in the ex vivo screening panel. The nominated agent was used as part of maintenance treatment for the primary high-grade glioma. The treatment decision was made independently of and without reference to the platform results.

For Pt7, WGS identified somatic *TSC1* loss, a candidate alteration in mTOR signaling (Figure 4D), and everolimus had an AUC below the 30th cohort percentile (Figure 4E), yielding a T1a assignment under the framework.

Three modalities implicated ALK as a candidate vulnerability. WGS identified an inversion involving ALK (Figure 4F), bulk RNA-seq detected an in-frame *ALK* fusion transcript (Figure 4G), and scRNA-seq showed *ALK* overexpression in the malignant-cell compartment (Figure 4H). These findings support expression of a structurally altered ALK locus but do not by themselves demonstrate constitutive signaling or drug dependence. The framework nominated lorlatinib as T2a because the molecular candidate was CNS-relevant and the drug was not included in the screen.

The *ALK* fusion was also detected by clinical testing, and the patient enrolled in PNOC008 (NCT03739372) and received lorlatinib as part of molecularly guided maintenance therapy. At the most recent follow-up, 3.5 years post treatment, relapse had not been observed. The platform’s independent nomination was therefore concordant with the administered agent, even if this single, multimodally treated case does not fully establish that lorlatinib caused the favorable course.

Concordance with clinical testing helps define the platform’s incremental information. UCSF500 detected the fusion at the DNA and transcript levels, whereas the study’s scRNA-seq data indicated that *ALK* expression was enriched in malignant cells. The framework placed lorlatinib alongside everolimus and other candidate nominations; it did not establish a comparative treatment ranking.

#### Subclonal resolution of therapeutic vulnerabilities

Intratumoral heterogeneity can contribute to treatment resistance and relapse^37,38^, as genetically and phenotypically diverse populations can undergo treatment-associated selection and clonal evolution^20–22^. Because the clinical interpretation of subclonal findings remains challenging, we evaluated whether the integrated data could support exploratory subclone-aware annotation.

Aggregate 120x WGS can support inference of variant clusters consistent with subclonal structure, subject to tumor purity, copy number, sampling, and sequencing assumptions. scRNA-seq can identify transcriptional clusters, and scBayes^35^ can associate single-cell profiles with WGS-defined assignments where variant coverage is sufficient. We therefore evaluated shared versus group-restricted expression candidates. These labels are exploratory and do not establish clonality, temporal ordering, or therapeutic susceptibility.

As a proof of concept, we applied this approach to Pt3, an ATRT-SHH tumor. Deep WGS supported two genomic subclone assignments with a proposed ancestor-descendant relationship (SC1 to SC2; Figure 5A). Unsupervised scRNA-seq analysis identified two transcriptional clusters, and scBayes preferentially assigned cells in the two clusters to SC1 and SC2, respectively (Figure 5B). Cluster-stratified analysis identified overexpression of *FGFR1*, *FGFR2*, *AXL*, *PDGFRB*, and *EPHA2* in both assignments (Figure 5C), generating hypotheses involving receptor-tyrosine-kinase signaling. SMO overexpression in both groups was consistent with the SHH classification. *ERBB2* overexpression was restricted to the SC2-assigned group (log2 fold change, +0.91), was not detected in SC1, and was absent from the pooled analysis. This pattern nominates a subclone-restricted *ERBB2*/HER2 hypothesis; and postulates sensitivity to lapatinib, tucatinib, or another HER2-directed agent.

**Figure 5.**
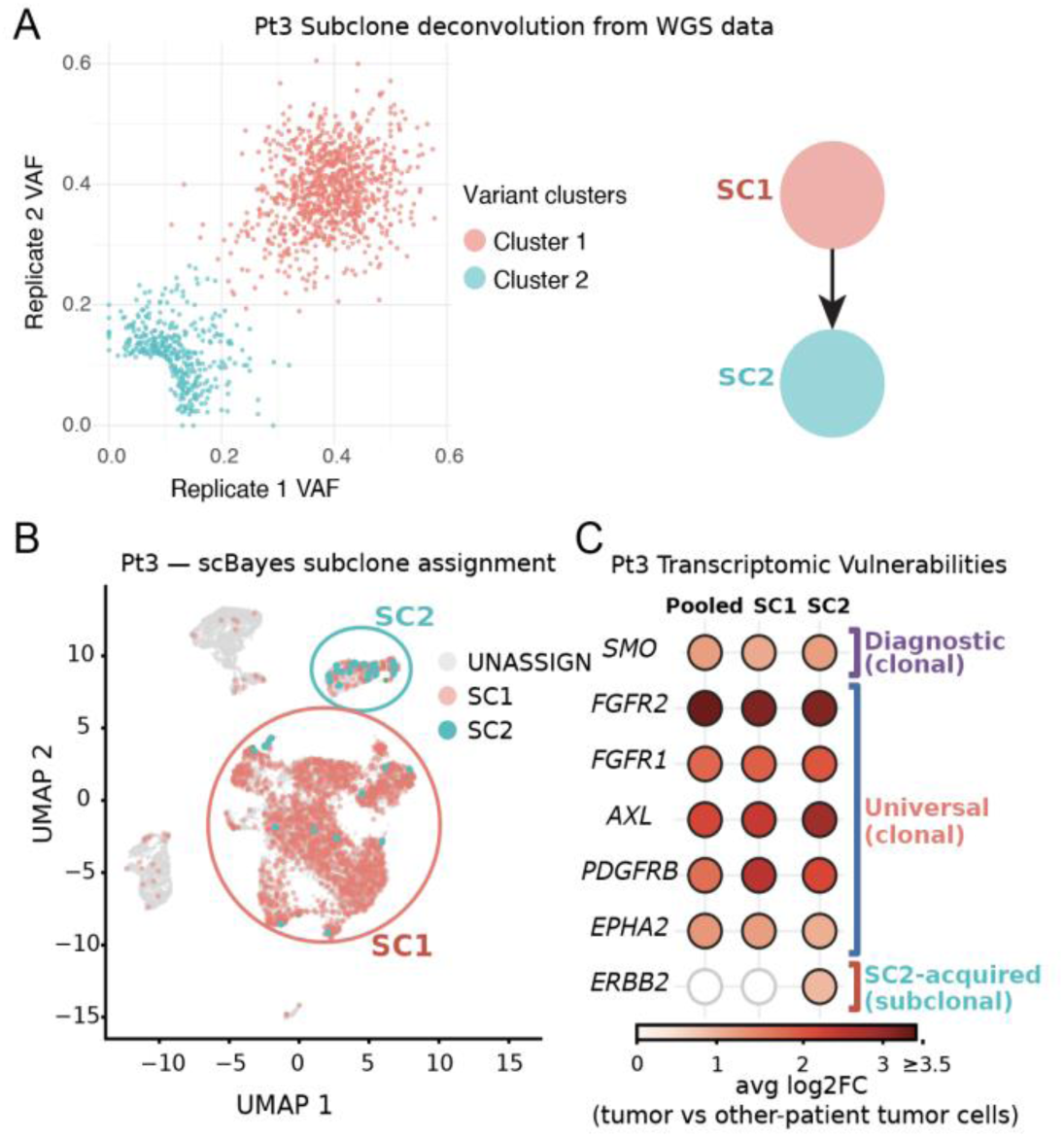
Subclonal resolution of therapeutic vulnerabilities in Patient 3 (ATRT-SHH). **(A)** Genomic subclone deconvolution from deep WGS. Somatic variant allele frequencies from two independent tumor replicates resolve two variant clusters corresponding to subclones SC1 and SC2 in an ancestor-descendant relationship (SC1 → SC2). **(B)** scBayes assignment of genomically defined subclones onto the scRNA-seq UMAP, mapping SC1 and SC2 to distinct transcriptional clusters (unassigned cells in gray). **(C)** Cluster-stratified transcriptomic vulnerabilities (average log2 fold change vs other-patient tumor cells) for pooled tumor cells, SC1, and SC2. *SMO* overexpression is universal and concordant with SHH classification (diagnostic/clonal); *FGFR2*, *FGFR1*, *AXL*, *PDGFRB*, and *EPHA2* are shared across both subclones (universal/clonal); *ERBB2* overexpression is acquired specifically in the descendant SC2 (subclonal), nominating HER2-directed therapy as a subclone-specific option.

We refined our tiered drug recommendation framework with specific annotations to indicate whether a given genomic or transcriptomic vulnerability is clonal or subclonal. These annotations will allow the treatment team e.g. to deliberately pair a trunk-targeting agent with subclone-targeting therapy - an approach particularly relevant at relapse, where resistant subclones may dominate the tumor mass. Pt3 therefore illustrates the feasibility of subclone-aware annotation in one tumor; systematic application and therapeutic evaluation remain future work.

## DISCUSSION

Precision oncology uses tumor molecular profiles to match patients with targeted therapies. In pediatric CNS tumors, current clinical practice includes targeted next-generation sequencing panels (e.g., UCSF500), DNA methylation classification, and RNA-based fusion testing. Because many pediatric brain tumors have few recurrent targetable alterations, these tests often yield limited matched-therapy options. Broader genomic profiling^9–11^, transcriptomic outlier analysis using bulk RNA sequencing^13,15^, and ex vivo functional drug testing^16,17^ have therefore been evaluated as complementary approaches.

This study evaluates three features of an integrated precision-oncology strategy (Tables 1 and 2). **First, the profiling layer combines sequencing depth (120x tumor WGS) with cellular resolution (scRNA-seq).** Deep WGS improves sensitivity for somatic mutation detection in specimens with substantial normal-cell admixture. At an aggregate depth of 120× and a 5% VAF calling threshold, clonal heterozygous variants remain detectable at tumor purities down to approximately 10-15%, and subclonal variants down to a cellular fraction of roughly 25% in a tumor of 40% purity - resolution that 30 - 60× sequencing does not afford. Across the 14 tumors with matched normal controls, FACETS estimated tumor purity ranged from 40% to 89%. scRNA-seq separated malignant-cell expression from stromal and immune signals, as illustrated for Pt3, Pt6, and Pt8 in Supplemental Figure 4, and nominated a reported mean of 37 and median of 42 druggable, cancer-associated overexpressed genes per patient. **Second, the inference layer adds an orthogonal functional axis and separates the logic for targeted and cytotoxic agents.** Track T ranked targeted agents by molecular-functional evidence, whereas Track C ranked six cytotoxic agents solely by cohort-relative ex vivo response. The resulting assignments are candidate nominations rather than demonstrated therapeutic options. **Third, the framework can resolve vulnerabilities at subclonal resolution.** In Pt3, linking WGS-defined subclones to scRNA-seq transcriptional clusters identified *ERBB2* overexpression confined to the descendant subclone and undetectable in the pooled analysis of the same tumor. These features generate candidate nominations rather than demonstrated therapeutic options, and the subclonal result establishes feasibility in a single tumor rather than general benefit.

This pilot study lays the foundation for prospective clinical evaluation of our platform. Our cohort spans five histologic diagnoses, demonstrating that the workflow, from tissue processing through tiered recommendations, generalizes across molecularly diverse CNS tumors. Beyond diagnostic breadth, clinical utility requires delivering recommendations within a relevant treatment decision window. In this study, our platform has returned tiered recommendations within approximately 3-4 weeks of surgical resection, with sequencing and drug screening proceeding in parallel. This timeline fits two clinical scenarios. For children with newly diagnosed high-grade glioma, the interval from surgery through radiation to the start of maintenance therapy is approximately six weeks, providing a comfortable window for platform-guided agent selection. For patients with relapsed disease, next-line therapy is typically initiated within three to four weeks of resection, and the platform can deliver recommendations within this window. We next plan a prospective Phase 0 feasibility study to evaluate whether this platform can generate integrated therapeutic recommendations within clinically relevant treatment windows. A larger, preferably diagnosis-stratified cohort will also support more stable expression references and drug-response rankings.

In conclusion, integrating genomic alterations, malignant-cell expression, and short-term ex vivo response generated candidate findings that no single modality recovered alone and made the contributing evidence explicit. The framework substantially expanded therapy nomination yield in this feasibility cohort, paving the way for future prospective validation studies to refine thresholds, examine reproducibility, and prove clinical utility. If validated, this approach can plausibly be applied to other cancer types in which clinical realities support advanced omic and functional screening characterization.

## METHODS

### Sample Collection and Processing

Fresh tumor tissue was collected at the time of surgical resection and transported to the laboratory within 24 hours for molecular and functional profiling. Tumor specimens were processed through two primary workflows. After verifying that sufficient tissue was obtained for pathologic diagnosis and any potential clinical trial enrollment, the remainder of the tumor was utilized for tumor banking. A portion of the tumor was snap-frozen in 1.8 mL Nunc freezer vials in a 70% isopropyl alcohol/dry ice slurry in the operating room for DNA and RNA extraction and subsequent sequencing. The remaining tissue made into viable single cells by mincing the tissue, followed by collagenase/dispace gentle enzymatic digestion, and 0.9 M sucrose gradient as previously described^39^ in detail in the Cheshier Laboratory. Dissociated tumor cells were cryopreserved in 1.8 mL Nunc freezer vials (10% DMSO/90% fetal bovine serum) either immediately (P0) or after short-term culture (P1) in serum free tumor growth media, as previously described^39^ and subsequently used for scRNA-seq and ex vivo drug screening. Matched peripheral blood samples were collected for germline sequencing in 14 patients; three patients underwent tumor-only sequencing because blood samples were not collected.

### Whole Genome Sequencing (WGS), Data Processing and Somatic Variant Calling

DNA was extracted from tumor tissue and matched blood samples using Qiagen DNeasy Blood & Tissue extraction kit (cat. no. 69504). At least 100 ng DNA per sample was used to make the sequencing library. Whole-genome sequencing libraries were prepared using NEBNext Ultra II DNA Library Prep Kit and sequenced on Illumina sequencing platform to achieve a target depth of 60× for each library. To ensure tumor sequencing quality, we prepared two libraries from each tumor sample and targeted in total 120×.

Raw sequencing reads were aligned to the human reference genome (GRCh38) using BWA-MEM version 0.7.17. Single-nucleotide variants (SNVs) and short insertions/deletions (indels) were identified using FreeBayes version 1.3.4. High quality variants were filtered by 1) variant quality > 20; 2) per sample sequencing depth > 15 and per tumor sample sequencing depth >30; 3) allele frequency (VAF) > 0.05 in at least one tumor library; 4) intersecting with 1000G genome accessibility mask of GRCh38; 5) filtering out multi-allelic variant sites. SNVs and short INDELs were annotated by SnpEff version 4.3. For tumors that have matched normal control, somatic variants were identified when the variant counts were 0 in the normal control sample. Three of the 17 patients in this cohort lack a matched normal sample. To recover somatic mutations for these tumors, we applied a filtering strategy that relies only on publicly available population resources and a cohort-derived Panel of Normals (PoN). The PoN was defined as the union of germline and artifact variants contributed by the 14 cohort patients with matched normal samples. Each patient’s contribution is the difference between their raw tumor-only call set and their matched-normal-derived somatic call set. For each tumor-only sample, raw variant calls were normalized with “bcftools norm”, applying multiallelic decomposition, left-alignment, and REF reconciliation against GRCh38. These variants were then filtered by progressive set subtraction with “bcftools isec”: variants present in gnomAD v4.1 or the 1000 Genomes Project were removed as likely germline, and the remaining variants were filtered against the PoN. The retained variants constitute the final tumor-only somatic call set.

Somatic copy number variants (CNV) and loss of heterozygosity (LOH) events were called by FACETS version 0.6.1, which uses the tumor to normal depth ratio to estimate copy number changes in the tumor and the ratio of alternative allele count to reference allele count (odds ratio) at single nucleotide polymorphism sites in the tumor sample to estimate LOH and allelic imbalance. For the three tumors without a matched normal control, we used a blood sample from an unrelated patient (Pt14) as a surrogate normal for copy-number calling. Because LOH detection requires a matched normal, it was not performed for these tumor-only cases.

Structural variants and break points were identified using the reference-free variant detection algorithm RUFUS d1.1.9, a reference-free k-mer-based detector^40^, followed by visual inspection in IGV^41^.

### Bulk RNA-seq, Data Processing and Fusion Detection

Total RNA was extracted using Qiagen RNeasy Plus mini extraction kits with DNase treatment (cat. No. 74134). RNA-seq libraries were prepared using NEBNext Ultra II Directional RNA Library Prep with poly(A) mRNA Isolation and sequenced on Illumina sequencing platform to generate 50 million paired-end reads per sample. Reads were aligned to the human reference genome (GRCh38) using STAR v2.7.11. Then count matrices were generated with featureCounts v2.1.1 and RSEM v1.3.3. Gene fusion detection was performed using STAR-Fusion v1.15.1.

### Single Cell RNA-seq and Data Processing

Single-cell RNA sequencing was performed on all 17 patient tumors. Cryopreserved single-cell suspensions were thawed and assessed for viability using a Countess II cell counter and Invitrogen Countess cell counting chambers with trypan blue exclusion (cat. no. C10283). Dead cells were removed by magnetic bead separation using EasySepTM Dead Cell Removal (Annexin V) kits (cat no. 17899). Samples with viability ≥ 30% were processed for scRNA-seq. Immediately prior to library generation, the cell viability was reassessed with a Cellometer K2 Fluorescent Cell Counter and the viable cell number was used to determine the amount loaded for our target cells per library. Single-cell libraries were prepared using the 10x Genomics Chromium platform 10X 3’ v3.1 with UDI according to the manufacturer’s protocol, targeting 6000 cells per sample. Libraries were sequenced on Illumina sequencing platform to achieve a minimum of 400 million read pairs. Raw sequencing data were processed using Cell Ranger version 7 with alignment to the human reference genome (GRCh38) and generate count matrices. ScRNAseq from multiple samples were aggregated using the CellRanger command “aggr” without depth normalization.

### Single-Cell Data Analysis and Cell Type Identification

Filtered count matrices were analyzed using Seurat v5. Quality control was performed to remove low-quality cells based on the following criteria: cells with fewer than 200 detected genes, cells with greater than 20% mitochondrial reads. After quality control, 962 to 16028 cells per sample were retained for downstream analysis (range: 316-12,495 tumor cells per sample). Data were normalized using SCTranform, and highly variable genes were identified for dimensionality reduction. Principal component analysis (PCA) was performed, and the top 30 principal components were used for uniform manifold approximation and projection (UMAP) visualization and graph-based clustering using nearest neighbor method. Cell type annotation was performed using known marker genes: *PTPRC -* immune cells; *MS4A1*-B cell; *NKG7,GNLY* - NK; *CD3E -* T; *CD14* - Monocyte; *CALCRL, RAMP2, PECAM1, CLDN5* - endothelial cell; *COL3A1, FBN1, LUM* - fibroblast cell; *MBP, TF, PLP1, MAG, MOG, CLDN11* - Oligodendrocyte (Supplemental Figure 1). Tumor cells were distinguished from normal stromal and immune populations based on somatic mutations and CNVs that tumor cells present (See Malignant-Cell Identification below).

We used the AggregateExpression function in Seurat to generate pseudobulk expression for scRNAseq data. For each sample, we applied this method to tumor cells only (pseudobulk for tumor cells) or to all cells (pseudobulk for all cells) (Supplemental Figure 4).

### Malignant-Cell Identification

Malignant cells were distinguished from non-malignant stromal and immune populations using a combined copy-number and genotype approach (Supplemental Figure 3B and C). First, inferCNV v1.3.3 was used to infer large-scale chromosomal copy-number alterations from single-cell expression profiles, using annotated immune and stromal cells as the diploid reference; cells exhibiting copy-number signatures concordant with the tumor’s bulk WGS copy-number profile were flagged as putative malignant. Second, scBayes was used to probabilistically assign individual cells to genomically defined tumor subclones on the basis of somatic variants identified from deep WGS, providing an orthogonal, genotype-based confirmation of malignant identity. Because the two methods rely on different signals, they compensate for each other’s limitations: for tumors lacking detectable copy-number alterations, malignant cells were identified using the somatic-mutation-based scBayes method (e.g. Pt1, Pt3, Pt9 etc., Supplemental Figure 3B), whereas for tumors without a matched normal control, malignant cells were identified using inferCNV (e.g. Pt6 and Pt17, Supplemental Figure 3C).

### Differential Expression Analysis

To mitigate sample-level and disease-level imbalance in downstream analyses, we downsampled to a maximum of 500 randomly selected tumor cells per patient. Patient-specific differential expression analysis was performed by comparing tumor cells from each patient to tumor cells from all other patients in the cohort (pan-CNS tumor reference). Differential expression was calculated using the non-parametric Wilcoxon rank-sum test with Bonferroni correction. Genes were considered significantly differentially expressed if they met the following criteria: adjusted p-value < 0.05 and log2 fold change > 0.5. Transcriptomic vulnerabilities were defined as significantly overexpressed genes that could be directly targeted by available drugs, as determined by annotation against curated drug-target databases (see Vulnerability Annotation).

### Cancer Gene Annotation

We constructed a cancer gene annotation system integrating six databases: the COSMIC Cancer Gene Census (758 genes)^25^, a pan-cancer driver gene set (633 genes)^26^, the TARGET database^27^, the Network of Cancer Genes (3,347 genes)^28^, OncoKB (1,203 genes)^29,30^, and the UCSF500 panel (480 genes) (Figure 1E, Supplemental Table 2). The union of these resources comprised 3,985 unique cancer-associated genes, each annotated for its role as an oncogene or tumor suppressor where available.

Somatic short variants (SNVs and indels), copy-number variants (CNVs) from WGS, and differentially expressed genes (DEGs) from scRNA-seq were annotated against the cancer gene system at the gene level. Short variants affecting any of the 3,985 cancer-associated genes were retained as candidate genomic vulnerabilities. CNVs were annotated with the additional requirement that the direction of the copy-number change be consistent with the gene’s oncogenic role: high-level amplification (copy number > 4) of an oncogene, or loss (heterozygous or homozygous deletion) of a tumor suppressor, was considered a vulnerability, whereas direction-discordant events (e.g., loss of an oncogene) were excluded. DEGs were likewise required to be cancer-relevant; their direction of dysregulation was incorporated at the drug-mapping stage.

### Multi-Source Drug Compendium

The therapeutic drug universe was assembled from four sources (in total 337 drugs): (1) a physician-curated, pediatric CNS tumor-relevant drug list (Supplemental Table 3); (2) the INFORM Drug Sensitivity Profiling panel^17^; (3) the Florida functional precision medicine panel^16^; and (4) all FDA-approved oncology drugs from OncoKB (Figure 1F). Twenty-five drugs were selected for ex vivo screening based on clinical relevance to pediatric CNS tumors and mechanistic coverage of key pathways, including RTK-MAPK, PI3K/AKT/mTOR, cell-cycle regulation, epigenetic modifiers, and cytotoxic agents (Supplemental Table 3).

### Drug-Target Knowledge Base and Vulnerability-to-Drug Mapping

For every drug in the compendium, we compiled its molecular target(s) and mechanism of action into a drug-target knowledge base integrating DrugBank, the Therapeutic Target Database (TTD), GDSC, and OncoKB, supplemented by manual curation, yielding a drug-vulnerability list that links each compound to the gene-level alteration(s) it is expected to engage. Genomic vulnerabilities (mutated genes and direction-matched CNV genes) were mapped directly to drugs whose targets included the altered gene. For transcriptomic vulnerabilities, mapping additionally required concordance between the direction of dysregulation and the drug’s mechanism of action: because the panel consists of target inhibitors, only overexpression of a drug’s target was treated as a targetable vulnerability (e.g., the WEE1 inhibitor adavosertib was nominated only when *WEE1* was overexpressed). Gene-drug pairs meeting these criteria defined the candidate targeted therapies carried forward to the two-track tiering framework (Supplemental Table 4).

### Ex-vivo Drug Screen and Data Analysis

The cryopreserved single cells were thawed at 37 °C for 5 min. Cells were seeded in a 384-well round-bottom low-attachment (spheroid) tissue culture plate (100-2000 cells per well) with 30 µl of Advanced DMEM/F12 media supplemented with 2% FBS. The plate was centrifuged at 400xg for 5 min and then incubated at 37 ℃ and 5% CO2 for 24 hours. Then 25 drugs were prepared with 5-point serial dilution and applied to the plate with 3 technical replicas. The final treatment doses were ranging from 0.01 to 32 µM. In addition, 9 wells were treated with the DMSO as the control. The plate with added drugs and DMSO was incubated for another 72 hours (37 ℃ and 5% CO2) and assayed with CellTiter-Glo.

Raw luminescent values from each condition were normalized to the value from the DMSO treated wells. Dose-response curves were fitted using a four-parameter log-logistic model, and drug efficacy was quantified using the area under the curve (AUC) which is estimated by sum of viability on the curve for each dose. For samples that cannot fit a dose-response curve, AUC was calculated based on the sum of viability of each dose. IC50 values were calculated where curve fitting permitted. To enable cross-patient comparison and account for differences in intrinsic drug potency, percentile scores were calculated for each drug by comparing its AUC in a given patient to the AUC distribution across the entire cohort. A drug at the 50th percentile reflects an average response relative to the cohort, while lower percentiles indicate greater sensitivity. Drugs with percentile <50 were defined as “effective” for a given patient (Supplemental Figure 5).

### Drug recommendation Tiering system

Candidate therapies were prioritized through a two-track framework reflecting the distinct inference logic for targeted therapy (Track T) and chemotherapy (Track C). In Track T, genomic alterations or transcriptomic outliers nominate drug-target hypotheses. When a nominated drug was included in the ex vivo panel, cohort-relative response supplied an orthogonal line of evidence. Drugs with a matched molecular candidate and an AUC percentile below 50 were assigned T1: T1a for percentile <30 and T1b for percentile 30-50. T1 therefore denotes molecular-functional concordance within this framework, not clinical or mechanistic validation. Molecularly nominated drugs not screened were assigned T2 and stratified as T2a (physician-curated pediatric CNS list), T2b (other FDA-approved oncology agents), or T2c (other investigational pediatric oncology agents). Screened targeted agents below the cohort median without a matched molecular candidate were assigned T3. A molecularly nominated drug with an AUC percentile ≥50 was flagged as discordant rather than labeled clinically resistant.

In Track C, the screen was the sole evidence source because the cytotoxic agents were not mapped to patient-specific molecular markers. Track C assignments were C1a (percentile <30), C1b (percentile 30-50), or Flagged-C (percentile ≥50). All percentiles are relative to the tested cohort and require prospective validation.

### Subclonal Reconstruction and Clonal-Aware Vulnerability Analysis

To resolve therapeutic vulnerabilities at subclonal resolution, we combined genomic subclone reconstruction from deep WGS with transcriptional clustering from scRNA-seq. Genomic subclones were reconstructed by clustering somatic variant allele frequencies across the two independent tumor replicates using SubcloneSeeker^42^, resolving distinct variant clusters and their ancestor-descendant relationships (e.g., SC1 → SC2 in Patient 3). Within the tumor cells in scRNAseq data, we used scBayes to probabilistically assign each single cell to a genomically defined subclone based on its observed somatic variants, linking transcriptional clusters to WGS-defined subclones where coverage permitted. When linkage was achievable, this produced a unified clonal map; when not, transcriptional clusters were retained as candidate subpopulations for vulnerability analysis. For each cluster or subclone, we performed independent differential expression against the pan-CNS tumor reference (as above) and annotated the resulting vulnerabilities as either universal (shared across all clusters) or subclonal (restricted to one). Subclonal vulnerabilities were explicitly flagged during drug nomination to support combination strategies pairing trunk-targeting and subclone-directed agents.

### Data and Code Availability

Processed single-cell count matrices are available at GSE348845. Targetable somatic mutations are listed in Supplemental Table 6.

## ACKNOWLEDGEMENT

The authors thank the patients who donated samples to this study. We thank all clinical personnel involved in the care of the patients. We thank the Utah Center for Genomic Medicine seed grant. X.H., Y.Q., G.S.B, C.R.S and G.T.M were partially supported by NIH grant U54CA224076, the Huntsman Cancer Institute Team Science Acceleration Award 24-TSA-01 and NCI award 3P30CA042014-34S5. Research reported in this publication used the High-Throughput Genomics and Bioinformatic Analysis Shared Resource at Huntsman Cancer Institute at the University of Utah and was supported by the NIH, NCI award P30 CA042014. We thank Opal Allen and Brian K. Dalley for their dedication for sequencing efforts. The computational resources used were partially funded by the NIH Shared Instrumentation Grant 1S10OD021644-01A1. We acknowledge the Molecular Discovery & Screening Core at the University of Utah for use of equipment (Tecan Evo96, HP D300 Dispenser, Bioteq Plate Reader), and thank Dr. Bai Luo for their assistance with these experiments. S.H.C was supported by donations and grants from Kathryn S. R. Lowry Endowed Chair In The Department Of Neurosurgery, Ty Louis Campbell Foundation, Intermountain Primary Children’s Hospital Center for Personalized Medicine, Intermountain Primary Children’s Hospital Foundation, Huntsman Cancer Foundation, Department of Neurosurgery-University of Utah, and Tate Family.

## Notes

### Competing Interest Statement

The authors have declared no competing interest.

